# Getting the Hologenome Concept Right: An Eco-Evolutionary Framework for Hosts and Their Microbiomes

**DOI:** 10.1101/038596

**Authors:** Kevin R. Theis, Nolwenn M. Dheilly, Jonathan L. Klassen, Robert M. Brucker, John F. Baines, Thomas C.G. Bosch, John F. Cryan, Scott F. Gilbert, Charles J. Goodnight, Elisabeth A. Lloyd, Jan Sapp, Philippe Vandenkoornhuyse, Ilana Zilber-Rosenberg, Eugene Rosenberg, Seth R. Bordenstein

## Abstract

Given the complexity of host-microbiota symbioses, scientists and philosophers are asking questions at new biological levels of hierarchical organization - What is a holobiont and hologenome? When should this vocabulary be applied? Are these concepts a null hypothesis for host-microbe systems or limited to a certain spectrum of symbiotic interactions such as host-microbial coevolution? Critical discourse is necessary in this nascent area, but productive discourse requires that skeptics and proponents use the same lexicon. For instance, critiquing the hologenome concept is not synonymous with critiquing coevolution, and arguing that an entity is not a primary unit of selection dismisses that the hologenome concept has always embraced multi-level selection. Holobionts and hologenomes are incontrovertible, multipartite entities that result from ecological, evolutionary and genetic processes at varying levels. They are not restricted to one special process but constitute a wider vocabulary and framework for host biology in light of the microbiome.

## Main Text

Holobiont is a term used to describe an individual host and its microbial community, including viruses and cellular microorganisms (1–6) (Figure 1). It is derived from the Greek word *holos* that means whole or entire. Microbial symbionts can be constant or inconstant, vertically or horizontally transmitted, and can interact with a host in a context-dependent manner as harmful, harmless or helpful. In most cases, the net outcome of these interspecies relationships varies with the presence of other symbionts. The term holobiont distinguishes itself by not only recognizing hosts and their obligate symbionts, but also emphasizing the diversity of facultative symbionts and their dynamic associations within a host. In contrast to binary host-microbial interactions, the properties of complex microbial communities and their hosts are newly appreciated and potentially universal. The host and microbial genomes of a holobiont are collectively defined as its hologenome (1, 2), and the pluralistic attributes of a holobiont scale directly to the hologenome (Figure 1). Microbial genomes can be stable or labile components of the hologenome, vertically or horizontally transmitted, and the functional traits that they encode are context dependent and may result in damage, benefit, or be of no consequence to the holobiont (7). Having settled on these terms, one can look at holobionts and hologenomes as incontrovertible realities of nature. Hologenome is fundamentally similar to the words genome and chromosome in that they reflect different levels of biological information. The terms holobiont and hologenome are therefore structural definitions, although their utility remains subject to debate (8).

**Figure 1.**
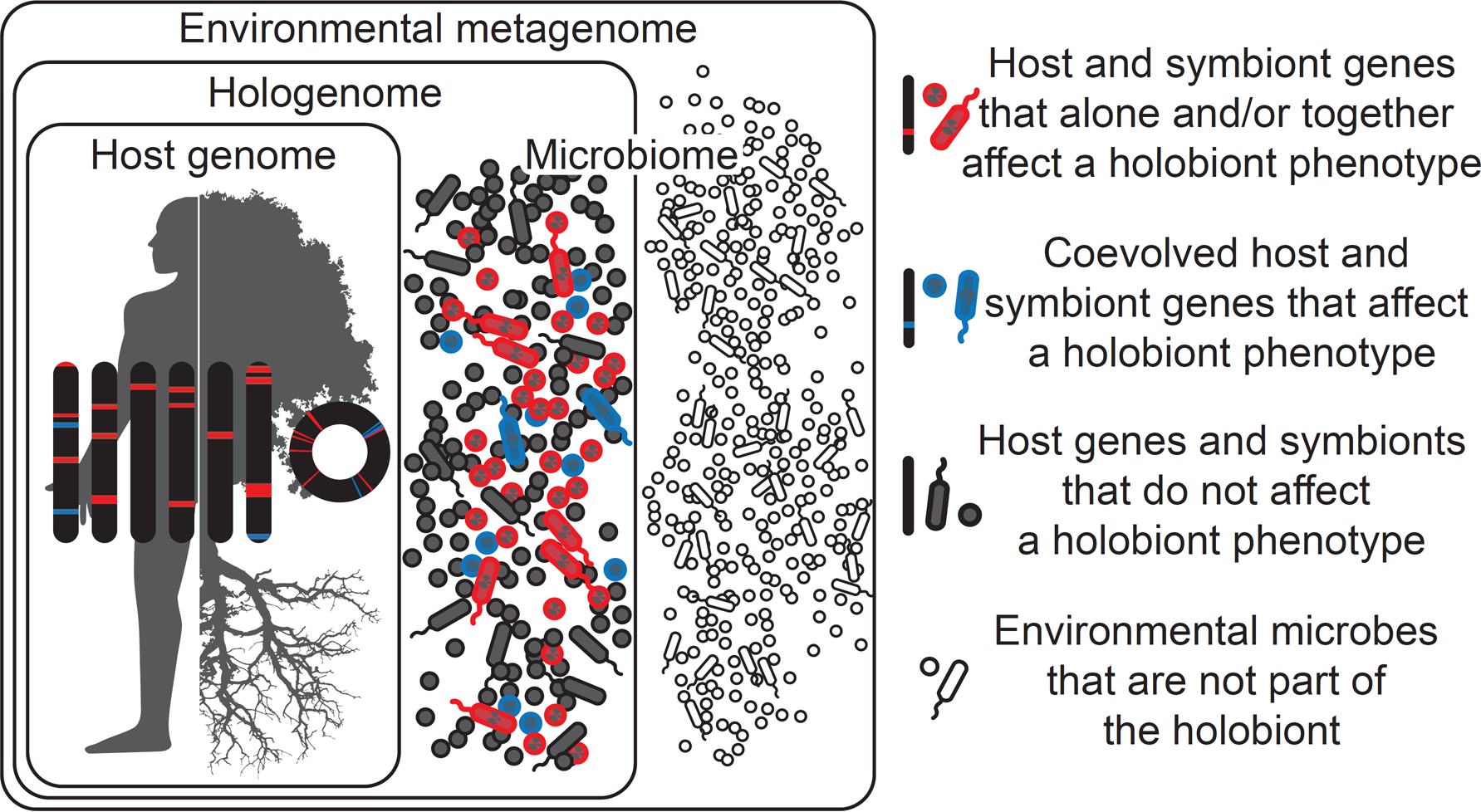
Holobionts are entities comprised of the host and all of its symbiotic microbes, including those which affect the holobiont's phenotype and have coevolved with the host (blue), those which affect the holobiont's phenotype but have not coevolved with the host (red), and those which do not affect the holobiont's phenotype at all (gray). Microbes may be transmitted vertically or horizontally, acquired from the environment, and can be constant or inconstant in the host. Therefore, holobiont phenotypes can change in time and space as microbes come into and out of the holobiont. Microbes in the environment are not part of the holobiont (white). Hologenomes then encompass the genomes of the host and all of its microbes at any given time point, with individual genomes and genes falling into the same three functional categories of blue, red and gray. Holobionts and hologenomes are entities, whereas coevolution or host-symbiont interactions are processes.

The first question then is why are these terms useful? They are useful because they replace misnomers in the context of host-microbiota symbioses like superorganism (i.e., an integrated social unit comprised of conspecifics), organ, and metagenome with a vocabulary that aligns with recent advances demonstrating that host phenotypes are profoundly affected by their complex microbial communities, in both cooperative and competitive ways (9–11). Holobionts and their hologenomes are less entities that elucidate something *per se* than they are entities that need elucidation.

The next question then is what is in need of elucidation? For any given symbiosis, host genetic variation may affect susceptibility to colonization of diverse microbes or even promote it in a highly specific way, microbial genetic variation may favor colonization while also affecting competition with coinfecting microbes, and environmental variation may substantively influence these dynamics and drive rapid microbial community changes. What is then in need of elucidation is how common and influential these forces are across host-microbial systems. Moreover, covariance between hosts and members of their microbiota is another important area of future research. Covariance can be achieved via vertical inheritance or selective filtering from the environment. The relative importance of these modes of holobiont assembly is not well resolved, yet either way, covariance in genetic compartments of the hologenome can yield variation in phenotypes upon which evolutionary processes can act. Finally, hologenomic variation may arise not only by mutation and recombination in the host and microbiome, but also by acquisition of new microbial strains from the environment, microbial amplification that involves a change in microbial abundance, and horizontal gene transfer among microbes (2).

Another area in need of elucidation is whether variation in traits caused by different host-microbiota assemblies drives a multigenerational response to selection. If there is a response to selection, then did it occur at the host, microbe, or microbial community levels? Can shifts at the microbial community level act akin to shifts in allele frequencies in host genomes? Preliminary indications are that not only can this occur, but that we can capitalize on its occurrence by artificially selecting (i.e., microbiome engineering) holobiont phenotypes in applied contexts (12, 13).

The hologenomic view of biology importantly does not prescribe host-centric or microbe-centric attributes to changes in holobiont macrobe functions, but rather takes into account the emergent interactions and outcomes of hosts and their microbiota. It is a relatively new view and is therefore liable to be interpreted in ways that misrepresent its original conception. For example, a recent paper expressed skepticism of the hologenome concept, yet did so by relying on alternative definitions that incorrectly restricted hologenomes to only those situations when holobionts are primary units of selection that arose by vertical inheritance and coevolution (8). The result is a straw man argument. The hologenome concept requires evaluation as any new idea does. However, to have a robust debate, skeptics and proponents must use the same terminology and framework. Here we highlight errors in these recent narrow definitions of the holobiont and hologenome, keeping them consistent with their original pluralistic definitions, and attempt to stimulate understanding of the link between holobiont phenotype and genotype.

The first argument proposed against the hologenome concept is that if X did not coevolve *sensu stricto* with Y, then the hologenome is not real (8, 14, 15). In this case, X and Y are respectively a microbe/microbial community and a host. As emphasized above and in the original literature, hologenome is a term that encompasses all of the genomes of the holobiont at a given point in time. Thus, holobionts can be formed through neutral processes, selection at the level of the host, symbiont or both (Figure 1). Although a component of it, coevolution is not the sole feature of the hologenome and its associated concepts. By way of illustration, one would not similarly say that if genes X and Y did not coevolve in a host, then they are not part of the same genome. Evolution of genomes and hologenomes is not a monolithic process, nor is it simply beanbag genetics. Genetic conflict, epistasis, selection, drift, etc. are all operational (1, 2). Thus, objections to the hologenome concept based on a lack of coevolution misrepresent what constitutes a hologenome and holobiont for that matter. To put it simply, coevolution is a process; the hologenome is an entity that embraces the eco-evolutionary processes inherent in much of macroscopic biology.

When referencing the original definitions of the hologenome, it was suggested that a non-coevolutionary application of the word hologenome would make it “sufficiently general that it can be interpreted in any number of ways” (8). This comment refers to the more generally accepted definition of the hologenome as all of the genomes in the holobiont, all of which in turn are evolving in that context (6). However, using the same logic, the word genome would be as unhelpful to biology as the term hologenome because it would be an insufficiently general definition of the types of evolutionary processes occurring within the genome. The main lesson here is that coevolution, genetic conflict, selection, and drift at multiple levels (host genomes, symbiont genomes, hologenomes) all occur. In arguing for a hologenomic status of macro-organisms, we noted that interspecies interactions underlying holobiont phenotypes follow a conceptual and theoretical continuum from genetic interactions or epistasis between genes in the same genome (1, 16). Both types of interactions can be transient or stable under varying conditions such as population structure and selection. Indeed, the (in)stability of a host and its horizontally-transmitted microbes follows a theoretical continuum under the same math from the (in)stability of interacting genes in the same genome that undergo recombination (16). Here we note that vertical transmission versus horizontal transmission is a false dichotomy to draw against the hologenome concept.

Prevalent misuse of coevolution in the microbiome literature is a legitimate concern and was the impetus for some of us coining the word “phylosymbiosis” (17). It describes the concordance between a host phylogeny (evolutionary relationships) and microbial community dendrogram (ecological relationships) based on the degree of shared taxonomy and/or abundance of members of the community (18–21). Phylosymbiosis does not *a priori* imply coevolution, cospeciation, cocladogenesis, or codiversification because this latter vocabulary implies concordant splitting of new species from a common ancestral one (19, 20, 22). Phylosymbiosis avoids these assumptions because it “does not presume that microbial communities are stable or even vertically transmitted from generation to generation” (19, 20). Rather, it refers to a pattern in which changes in separate parts of the holobiont (host and microbiota) are related in a concordant manner. It is also a stepping-stone from population genetics to community genetics because when phylosymbiosis is observed under strictly controlled conditions, it tests whether variation in holobiont assembly is primarily stochastic or deterministic (17, 19, 23). Stochastic assembly means that each microbe has an equal opportunity of colonizing a host. Deterministic assembly reflects ecological selection of a particular non-random microbial community and its host, without reference to which partner, or potentially both, is doing the selecting, and it can be affected by genetic variation in the host or microbial species. Controlled studies of microbial community assembly across different species of *Nasonia* wasps and *Hydra* have yielded such phylosymbiotic patterns (17, 18). When genetic variation in the interacting species affects community assembly, it has been defined as broad sense “community heritability,” or *H^2^_c_* (24, 25). Similar to population genetic heritability estimates of phenotypes that are abiotic, *H^2^_c_* measures a “heritable basis to trophic-level interactions” (26). If there is a significant *H^2^_c_*, natural selection can act on genetic variation affecting ecological community structure (23, 27), including organization of the holobiont and its emergent phenotypes (25).

Discussion of evolutionary processes brings forth a second argument against the hologenome concept, namely that holobionts and their hologenomes must be the “primary” unit of selection (8). This strict claim leads biologists into error, as all of the literature emphasizes the reality that multiple levels of selection can operate simultaneously. For example, selfish genetic elements can be selected within a genome that is in turn selected for any number of phenotypes that affect fitness—this is uncontroversial. While the holobiont is posited to be “a unit of selection in evolution” (2, 28–30), it is naturally not proposed as the only or necessarily primary unit of selection (1, 2). Primariness varies with what traits are targeted by natural selection.

As we have emphasized in different venues, it is also true that just as large parts of the nuclear genome can evolve neutrally or be in conflict, so too can large parts of the hologenome (1, 2). For example, “hologenomic drift can occur at all the different levels of the holobiont from single genes of the microbes or the host to the holobiont itself” (2). We would be remiss to not be critical of our own inconsistent statements about the relative roles of cooperation and conflict in hologenomic evolution. In *The Hologenome Concept*, some of us stated that “evolution of animals and plants was driven primarily by natural selection for cooperation between and with microorganisms” (2) while in other venues the concept “places as much emphasis on cooperation as on competition” (31). This latter statement is more precisely aligned with the pluralistic nature of the holobiont, namely that “natural selection…on holobiont phenotypes…can work to remove deleterious nuclear mutations or microbes while spreading advantageous nuclear mutations or microbes” (1). In fact, some of us argued that conflicts of interests resulting from the nature of the transmission of microbes to the next host could select for microbes that can manipulate the biology of their host to improve their own transmission (32). The holobiont is not a conglomerate that arises solely from cooperation. Rather, it is a hierarchical level that can supersede the individual host that lives in association with its microbial community, incorporating both competitive and cooperative selective systems (33). Hologenomes then exist as hierarchically nested, although not necessarily integrated levels of genomes, in which all levels of selection are in play.

In summary, we anticipate that many subdisciplines in biology will benefit from a conceptual, theoretical, and experimental framework that broadly encompasses the ecology of holobionts and evolution of hologenomes. The hologenome concept is a comprehensive and relevant eco-evolutionary framework for which critical questions remain. For example, can a response to selection on host traits be driven solely by changes in the genomes and/or membership of a host-associated microbial community? How taxonomically widespread among hosts is phylosymbiosis? How common is vertical inheritance of complex microbial communities? What is the strength of selection required to maintain consistent association between a host and environmentally-acquired microbes each generation? How does selection operate on community phenotypes if *H^2^_c_* is variable due to the lability of microbial communities? Evolution of the hologenome refers to the genetic basis of eco-evolutionary processes underlying community phenotypes of the holobiont. This terminology and framework for the newly appreciated complexities in the host-microbe consortia and their genomes is not restricted to one special process but constitute an incontrovertible vocabulary and framework for host biology in light of the microbiome.

## Acknowledgements

We thank Jay Evans and Phil Pellett for helpful feedback on the manuscript.

